# The mineralization characteristics of organic carbon and particle composition analysis in reconstructed soil with different proportions of soft rock and sand

**DOI:** 10.1101/636993

**Authors:** Zhen Guo, Jichang Han, Yan Xu, Chang Tian, Chendi Shi, Lei Ge, Juan Li, Tingting Cao

**Author notes:** Corresponding author /.

## Abstract

The organic carbon mineralization process can reflect the release intensity of soil CO_2_. Therefore, the study of organic carbon mineralization and particle composition analysis of soft rock and sand compound soil can provide technical support and theoretical basis for the theory of soil organic reconstruction. Based on the previous research, this paper mainly selected four typical treatments of 0:1 (CK), 1:5 (C1), 1:2 (C2) and 1:1 (C3), respectively, and analyzed the soil organic carbon mineralization process and particle composition by lye absorption method, laser particle size meter and scanning electron microscope. The results showed that there was no significant difference in organic carbon content between C1, C2 and C3 treatments, but they were significantly higher than CK treatment (P < 0.05). The organic carbon mineralization rate of each treatment accords with a logarithmic function throughout the culture period (P < 0.01), which can be divided into a rapid decline phase of 1-11 days and a steady decline phase of 11-30 days. The cumulative mineralization amount on the 11th day reached 54.96%-74.44% of the total mineralization amount. At the end of the culture, the cumulative mineralization and potential mineralizable organic carbon content of C1 and C2 treatments were significantly higher than those of CK treatment, and the cumulative mineralization rate was also the lowest with C1 and C2 treatment. The turnover rate constant of soil organic carbon in each treatment was significantly lower than that of CK treatment, and the residence time increased. With the increase of volume fraction of soft rock, the content of silt and clay particles increases gradually, the texture of soil changes from sandy soil to sandy loam, loam and silty loam, and because of the increase of small particles, the structure of soil appears to collapse when the volume ratio of soft rock was 50%. In summary, the ratio of soft rock to sand volume was 1:5-1: 2, which can effectively increased the accumulation of soil organic carbon. At this time, the distribution of soil particles was more uniform, the soil structure was stable, and the mineralization level of unit organic carbon was lower. The research results have practical significance for the large area popularization of soft rock and sand compound technology.

## Introduction

The farmland soil organic carbon pool plays an important role not only in the process of global carbon circulation but also as the most important material base for soil fertility [1], and has a decisive role in the maintenance of cultivated land productivity, the prevention and treatment of soil erosion, the spatial and temporal variation of soil respiration and its stability, etc [2]. The soil organic carbon pool in terrestrial ecosystems is about 3 times that of plant carbon pools, and the organic carbon exchanged between soil and atmosphere accounts for about 2/3 of the total carbon storage of surface ecosystems, which slightiy changeed will have a certain impact on greenhouse gas emissions [3]. Soil organic carbon mineralization is an important part of the carbon cycle in terrestrial ecosystems. The change of land resources caused by human activities often cause changes in atmospheric CO_2_ concentration through the effects on terrestrial ecosystems, which in turn affects the carbon cycle and the climate change process [4]. Therefore, in recent years, the research on soil carbon cycle by human activities has attracted much attention and has become the core issue of multidisciplinary research.

Soil organic carbon mineralization is a process in which organic substances are decomposed into inorganic substances under the action of microbial degradation, providing nutrient content for crop growth and releasing greenhouse gases such as CO_2_ to the outside world [5]. Currently, studies have shown that arid and semi-arid regions account for 41% of the global land area, carrying 38% of the population and are sensitive to global climate change and human activities [6]. In the arid and semi-arid areas of Shaanxi, Shanxi, Inner Mongolia and other arid areas, as two important soil resources, the soft rock weathering (soft rock) and aeolian sandy soil (sand) are serious soil erosion and loose texture, and the nutrient content is low and the structure is poor. Chinese and foreign experts called it “Earth Environment Cancer”, it can be seen that the geographical hazard of the soft rock and sand areas determines the urgency and difficulty of ecological restoration in the region [7]. Han et al. [8] studied the structure and physicochemical properties of sandstone and sand, and analyzed that the two resources can be mixed into different proportions to form “new soil”, and also suggested that the optimum proportion of crops suitable for planting was range between 1:5 and 1:1 (soft rock: sand). At present, the technology of using soft rock to improve aeolian sandy soil has been widely used, and 1573.3 hm^2^ of newly cultivated land has been added, which has realized the resource utilization of the soft rock and the improvement of the regional ecological environment. She et al. [9] studied the nutrient content and hydraulic parameters of the soft rock and sand compound soil, and proposed that the fertilizer retention performance of the compound soil can be effectively improved with the increase of the soft rock content, and the available nutrients content can be improved when the water was sufficient. Wang [10] mixed sand and soft rock according to a certain proportion to form a new type of soil, mainly analyzed its texture changes and biochemical indexes of crops, and considered that the soil texture changed from sand to silt loam as the proportion of soft rock increases. Moreover, the photosynthesis rate of crops and the activity indexes such as antioxidant enzymes showed a trend of increasing first and then decreasing. It can be seen that for the two low-utilization resources of soft rock and aeolian sandy soil, many scholars have already carried on the resource utilization and obtained the certain achievement, and the research report on the combination effect of the soft rock and the aeolian sandy soil has gradually become more and more. However, under the background of global warming at present, we should not only improve the utilization rate of waste resources but also maintain the sustainable development of ecological environment. The issue of greenhouse gas emissions from the compounded soil is an area which has not yet been involved by researchers at present. The carbon source or sink of the composite soil also needs to be further studied.

Due to the large amount of montmorillonite in the soft rock and its strong hydrophilicity and adsorption, while the aeolian sandy soil is leaking water and leaking fertilizer, the texture is loose, which is complementary to each other in nature [11]. The team members have done some research on the soft rock and sand compound soil in the early stage. The results showed that the increase of the ratio of soft rock to sand can effectively increased the capillary porosity and decrease the water infiltration coefficient of the aeolian sandy soil. The texture was also improved to a certain extent, and the macroscopic mechanics showed strain hardening phenomenon and nonlinear characteristics. After adding soft rock, the final water content of the improved soil was significantly higher than that of the aeolian sandy soil, which was beneficial to the maintenance of water and fertilizer [11–14]. The results of team study also showed that the toxic effect of lead in aeolian sandy soil can be effectively reduced with the addition of the proportion of soft rock, and the larger the proportion of soft rock added, the greater the degree of reduction [13]. It can be seen that the research team mainly studied the hydraulic properties, fertility and adsorption of the soft rock and aeolian sandy soil in the early stage, but lacked the research on the carbon source carbon sink effect of the compound soil. Therefore, the purpose of this study were to (1) clarify the carbon mineralization strength of the compound soil in different proportions of sandstone and sand; (2) understand the microstructure and particle composition of the compound soil; and (3) clarify the carbon fixation effect of different proportion of mixed soil and provide the way and basis for the sustainable development of regional ecology. We hypothesized that (1) the addition of different proportions of soft rock can effectively increase the soil organic carbon and mineralized carbon content, and can play the role of carbon sink when the volume fraction of soft rock is less than 50%; and (2) with the increase of volume fraction of soft rock, the soil fine particles increase, and the texture changes from sand to silt loam.

## Materials and methods

### Overview of the test site

The experimental plot was set up in Fping County pilot test base of Shaanxi provincial land engineering technology research institute. Fuping County (108°57′-109°26′E, 34°42′-35°06′N) is the transition zone between the Guanzhong plain and the northern Shaanxi plateau, and belongs to the gully region of the Weibei Loess Plateau. The terrain is high in the north and low in the south. It slopes from the northwest to the southeast. The elevation in the territory is 375.8-1420.7 m. The area belongs to the continental monsoon warm zone with semi-arid climate. The annual total radiation is 5187.4 MJ m^-2^, the annual average sunshine hours is about 2389.6 h, the annual average temperature is 13.1 °C, and the annual average precipitation is 527.2 mm. The interannual variation of precipitation is large, and the annual precipitation coefficient of variation (CV) reaches 21.1%.

### Experiment design

The field test plot was to simulate the land condition of the soft rock and sand mixed layer in the Mu Us sandy land. The experimental plot was to lay a mixture of soft rock and sand at 0-30 cm, and fill the aeolian sandy soil with 30-70 cm. The soft rock and sand were taken from Daji Han Village, Xiaoji Han township, Yuyang district, Yulin city. The minerals in the soft rock mainly include quartz, montmorillonite, feldspar, calcite, illite, kaolinite and dolomite. The main chemical constituents of soft rock are SiO_2_ (65% by mass), Al_2_O_3_ (14% by mass), Fe_2_O_3_ and CaO (21% by mass). The mineral in the sand is mainly quartz (SiO_2_), and its mass fraction is about 82%. The remaining minerals are mainly feldspar (10% by mass), kaolinite (4% by mass), calcite (2% by mass) and amphibole (2% by mass).

The test was carried out in 2009, and four treatments of soft rock and sand in a volume ratio of 0:1 (CK), 1:5 (C1), 1:2 (C2), and 1:1 (C3) were selected. Each treatment was repeated 3 times for a total of 12 trial plots. The area of the plot was 2 m×2 m = 4 m^2^. According to the site conditions of the test plot, considering the uniformity of factors such as illumination and micro-topography, the test plot was arranged from south to north in a “one” shape, usually the depth of the soil layer was 30-40 cm, so the mixing depth of the soft rock and sand in the test plot was designed to be 0-30 cm, simulating the field conditions, and 30-70 cm was completely filled with sand. The experimental field was corn (Jincheng 508)-wheat (Xiaoyan 22), which was made by two crops a year, all of which were artificially sown. The types of fertilizers tested in the experimental field were urea (including N 46.4%), diammonium phosphate (including N 16%, containing P_2_O_5_ 44%), potassium sulfate (including K_2_O 52%), and the amount of fertilizer applied was 255 kg hm^-2^ (N), 180 kg hm^-2^ (P_2_O_5_) and 90 kg hm^-2^ (K_2_O). All treatments were fertilized the same way. All phosphate fertilizers and potassium fertilizers were used as base fertilizers, 65% of nitrogen fertilizers are used as base fertilizers, and 35% were combined with irrigation water to be applied at the booting stage. Wheat planting time is generally in the middle and late October, and corn planting time is in the middle and late May. One to two days before sowing, the three fertilizers should be weighed according to the required amount of each plot, and evenly spread on the soil surface, and then properly ploughed to mix the fertilizer with the topsoil.

### Soil sample collection

After the wheat harvest in May 2018, samples of 0-30 cm soil layers in each plot were collected, and each plot was uniformly collected 5 points to form a mixed sample. The collected soil samples were removed from animals and plant residues, sieved through a 2 mm sieve and divided into two parts, one part was placed in a 4 °C refrigerator for mineralization culture test; one part was naturally air-dried and ground 1 mm sieve and 0.149 mm screen for scanning electron microscopy analysis and determination of organic carbon and texture.

### Determination method

The mineralization culture test was carried out by the alkali absorption method [15]. Organic carbon was determined by potassium dichromate-concentrated sulfuric acid external heating method [16]. The texture was measured using a Malvern laser particle size analyzer (MS2000, UK). The soil sample after 1 mm screen was cured by epoxy resin, and coarsely ground, artificially ground and polished by a sander to make the surface smooth and smooth, and a micro sample with a diameter of 5 mm and a height of 3 mm was obtained. The dried sample was subjected to gold plating, and scanning was performed by a scanning electron microscope (FEI Q45, USA) having a voltage of 15 kV in an “S” type, and the magnification was 1000 times.

### Data Processing and Analysis

Texture data was classified using TriangleVB software. All data were sorted and graphed using EXCEL 2019, and analysis of variance and multiple comparisons were performed using SPSS 19.0.

## Results

### Compound soil organic carbon

The organic carbon content of aeolian sandy soil can be significantly improved by adding different proportion of soft rock (**Fig 1 and S1 Table**). The organic carbon content of CK treatment was 2.02 g kg^-1^, and the organic carbon content of C1, C2 and C3 treatment was significantly increased compared with that of CK treatment (P < 0.05), with increases of 110.24%, 76.72% and 118.77%, respectively. With the increase of the proportion of soft rock, the soil organic carbon increased first and then decreased, then continued to increase. The organic carbon content of C1 treatment was 4.24 g kg^-1^, and the organic carbon content of C2 and C3 treatment decreased and increased by 15.94% and 4.06%, respectively, and there was no significant difference (P > 0.05). The C3 treatment increased by 23.80% compared with the C2 treatment, and there was no significant difference.

**Figure 1.**
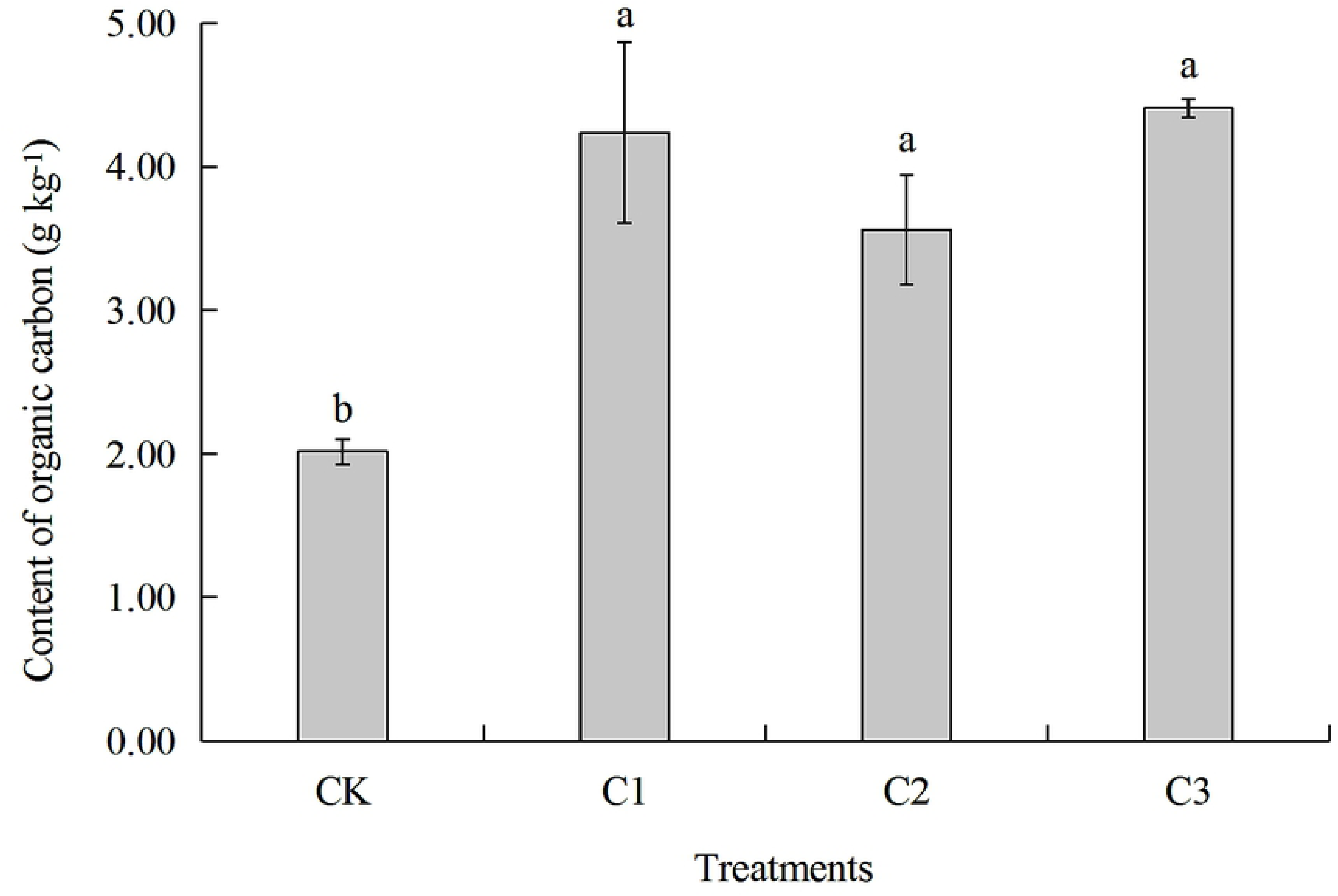
Organic carbon content of compound soils in different proportions of soft rock and sand. Different letters above the bars mean significant difference (at 0.05 level) between treatments. CK: the volume ratio of soft rock to sand is 0:1; C1: the volume ratio of soft rock to sand is 1:5; C2: the volume ratio of soft rock to sand is 1:2; C3: the volume ratio of soft rock to sand is 1:1.

### Compound soil organic carbon mineralization rate

The mineralization rate of organic carbon in soils with different mixing ratios of soft rock and sand showed a dynamic downward trend with the cultivation time (**Fig 2 and S1 Table**), which conforms to the logarithmic function relation y = a + b ln(x) (P < 0.01) (**Table 1**). The mineralization rates of organic carbon in CK and C3 treatments reached the peak on the 3rd day of incubation time, was 49.61 mg kg^-1^ d^-1^ and 66.07 mg kg^-1^ d^-1^, respectively, which increased by 76.48% and 37.19% compared with the first day. The mineralization rate of organic carbon decreased rapidly after 3rd day and began to decline slowly after 11th day. The organic carbon mineralization rate at the 30th day of culture was 3.41 mg kg^-1^ d^-1^ (CK) and 13.48 mg kg^-1^ d^-1^ (C3), respectively, which decreased by 93.13% and 79.60% compared with the 3rd day. However, the organic carbon mineralization rate of C1 treatment and C2 treatment reached the maximum on the first day of culture, which was 58.67 mg kg^-1^ d-^1^ and 43.51 mg kg^-1^ d^-1^, respectively. The mineralization rate of C1 and C 2 treatments were in a rapid decline stage until the 11th day, after that there was a slow descent. The mineralization rate on the 30th day was 82.65% and 81.68% lower than that on the first day, respectively. On the whole, the average mineralization rate of C3 treatment was the highest in all compound ratio treatments, followed by C1 treatment and C2 treatment, and CK treatment had the lowest mineralization rate. The mineralization rate of all treatments can be divided into two stages of change, namely rapid decline (1-11 days) and steady decline (11-30 days). The CO_2_ production rate changes greatly during the rapid decline phase, and the mineralization rate between all treatments tends to be consistent during the steady decline phase.

**Table 1.**
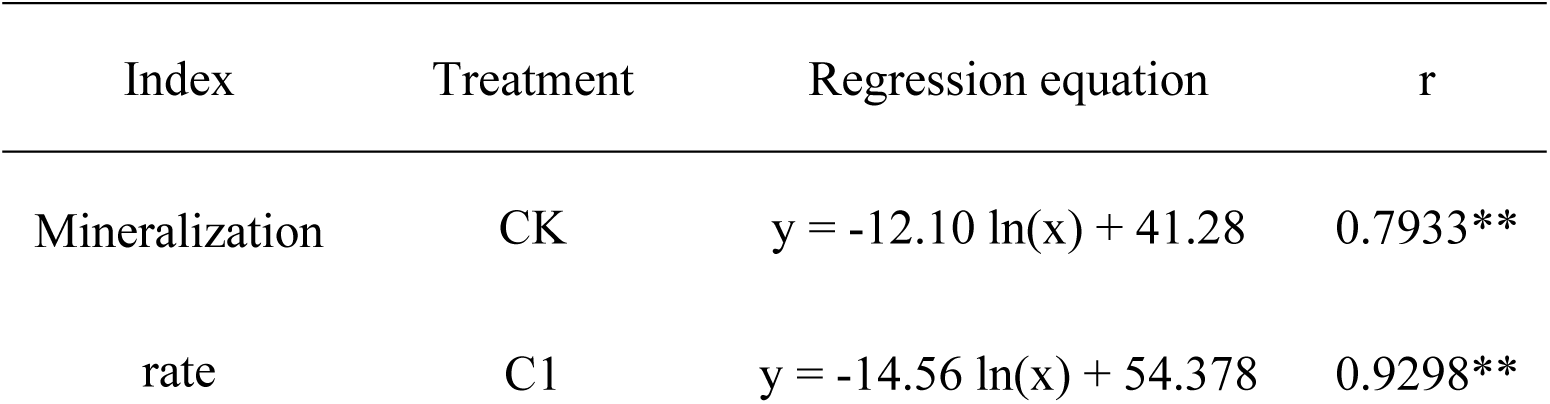
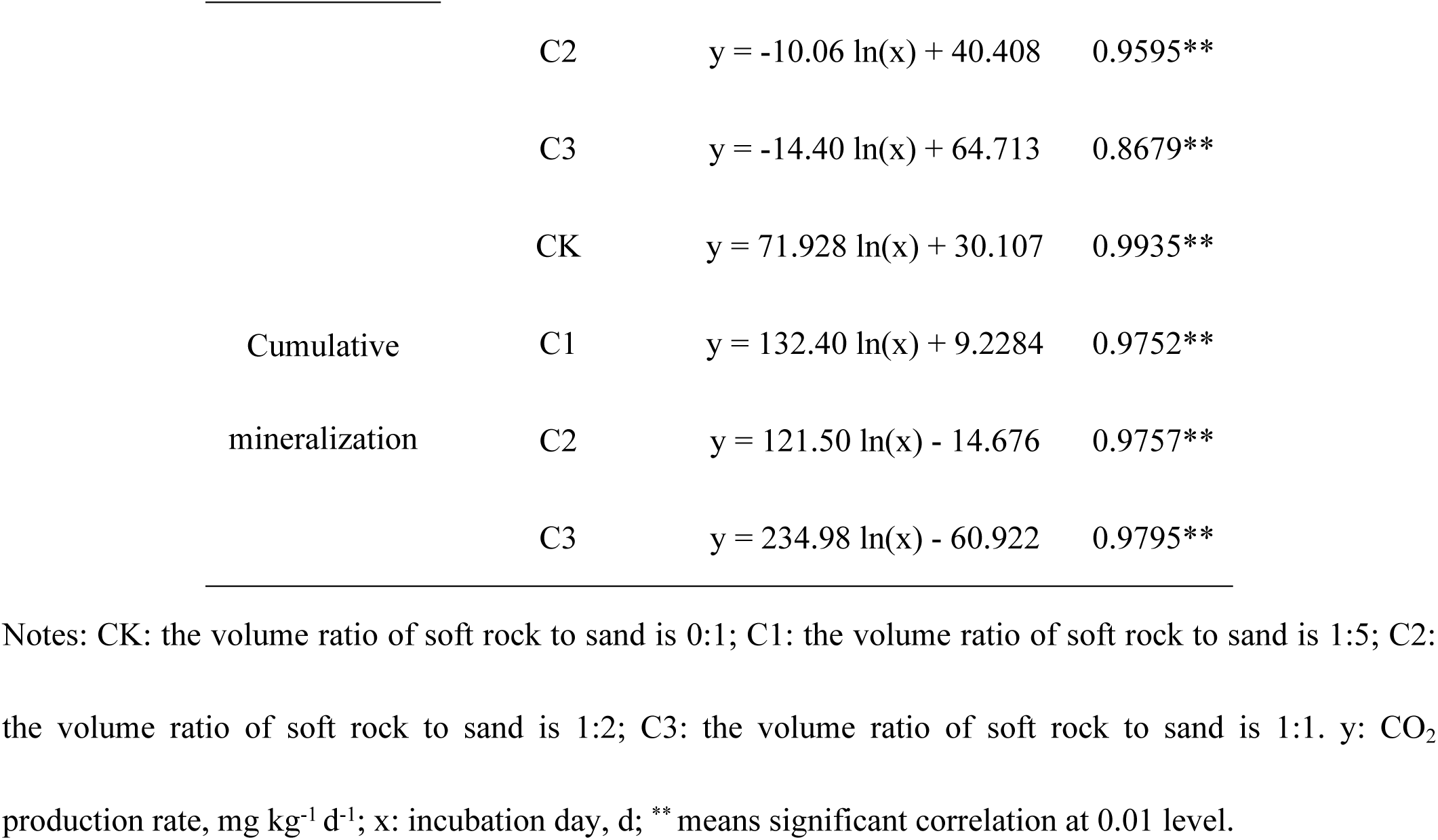
Regression equation of soil organic carbon mineralization rate and cumulative mineralization under different compounding ratios.

**Figure 2.**
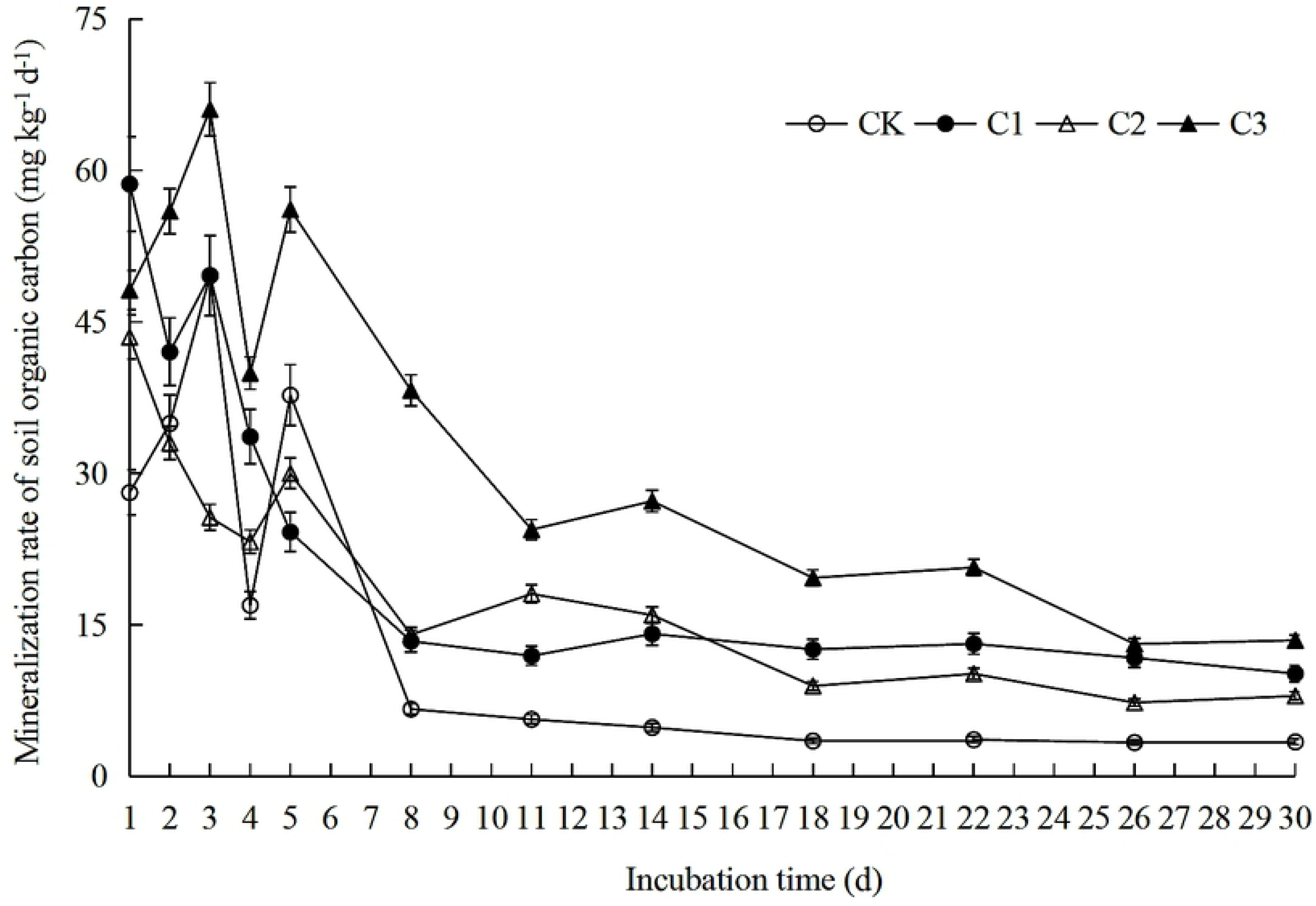
Organic carbon mineralization rate of compound soils in different proportions of soft rock and sand. CK: the volume ratio of soft rock to sand is 0:1; C1: the volume ratio of soft rock to sand is 1:5; C2: the volume ratio of soft rock to sand is 1:2; C3: the volume ratio of soft rock to sand is 1:1.

### Cumulative mineralization of compound soil organic carbon

The relationship between the cumulative mineralization of organic carbon and the incubation time in different proportions of soft rock and sand was also in accordance with the logarithmic function relationship y = a + b ln(x) (P < 0.01) (**Fig 3 and Table 1**). The results showed that the accumulation intensity of organic carbon accumulation mineralization decreased gradually with the extension of incubation time, that was, the CO_2_ release rate decreased. During the whole culture period, the cumulative mineralization of organic carbon in C3 treatment was the highest, followed by C1 and C2 treatments, and CK treatment had the lowest accumulation mineralization, with significant difference among all treatments (F = 26.54, P < 0.01). After incubation for 30 days, the cumulative mineralization of organic carbon treated by CK was 274.44 mg kg^-1^. The cumulative mineralization of organic carbon treated by C1, C2 and C5 increased by 88.40%, 59.32% and 192.92%, respectively, and the results reached significant differences (**Table 2**). There was no significant difference in the cumulative mineralization of organic carbon between the C1 and C2 treatments. Compared with C1 treatment and C2 treatment, the cumulative mineralization of organic carbon in C3 treatment was significantly increased by 55.48% and 83.86%, respectively.

**Table 2.** Cumulative mineralization of SOC after the 30 days of incubation and parameters of its kinetic equations.

| Treatment | $C_t$(mg kg<sup>-1</sup>) | $C_0$(mg kg<sup>-1</sup>) | k (d<sup>-1</sup>) | $T_{1/2}$ | $C_0$/SOC (%) | $R^2$ |
| --- | --- | --- | --- | --- | --- | --- |
| CK | 274.44 c | 257.44 c | 0.1671 a | 4.15 b | 12.78 b | 0.9748** |
| C1 | 517.03 b | 526.05 b | 0.0824 b | 8.41 a | 12.42 b | 0.9643** |
| C2 | 437.22 b | 484.61 b | 0.0697 b | 9.94 a | 13.61 b | 0.9947** |
| C3 | 803.88 a | 936.95 a | 0.0622 b | 11.14 a | 21.25 a | 0.9983** |

Notes: CK: the volume ratio of soft rock to sand is 0:1; C1: the volume ratio of soft rock to sand is 1:5; C2: the volume ratio of soft rock to sand is 1:2; C3: the volume ratio of soft rock to sand is 1:1. $C_t$ for amount of organic carbon cumulative mineralization, $C_0$ for amount of potential mineralizable organic carbon, k for constant of organic carbon mineralization rate, $T_{1/2}$ for half turnover period, $C_0$ /SOC for ratio of potential mineralizable organic carbon to total organic carbon in compound soil. Values followed by different letters in the same column mean significant difference at 0.05 level between treatments, \*\* indicates a extremely significant level of 1%.

**Figure 3.**
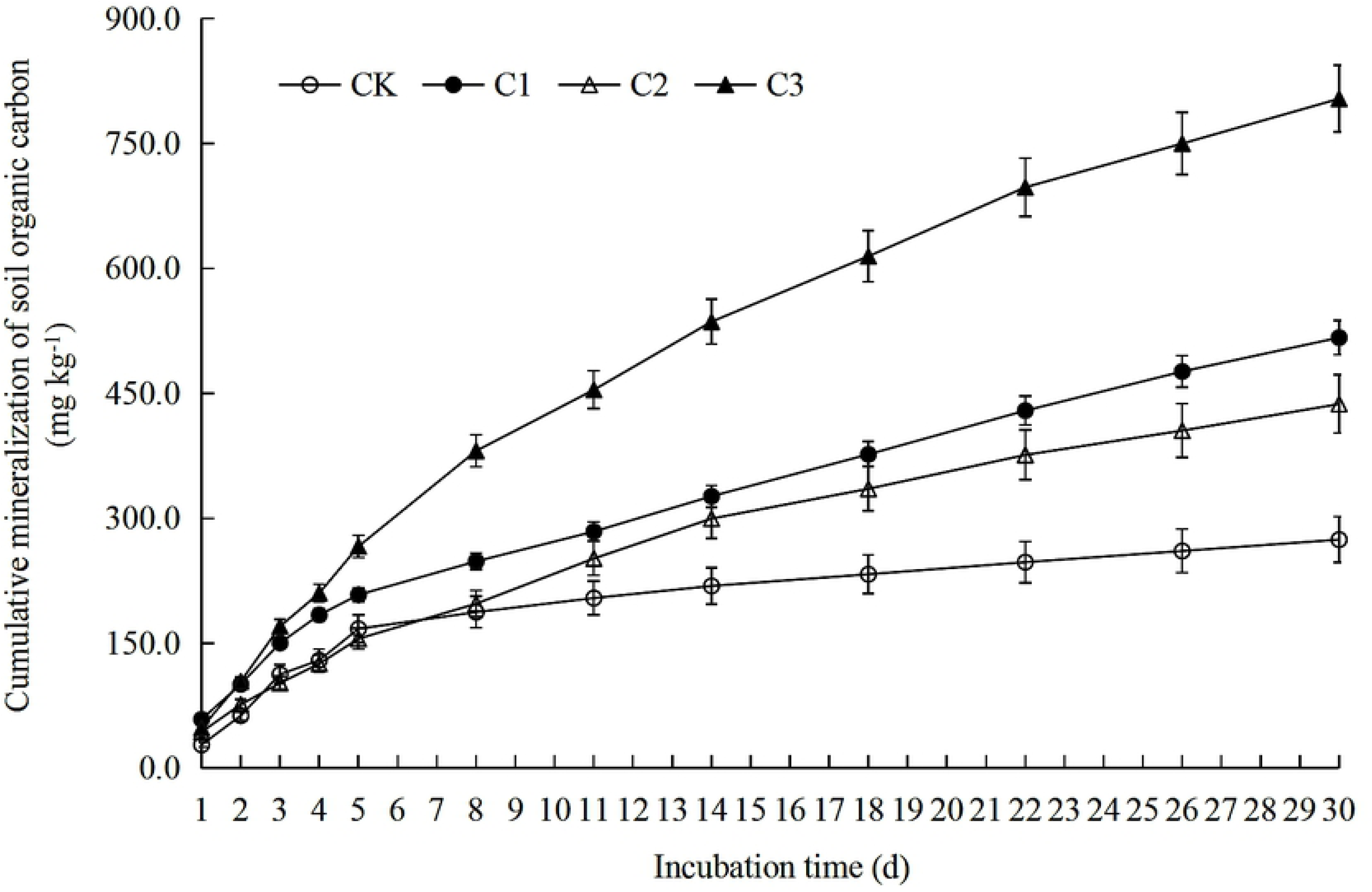
Organic carbon cumulative mineralization of compound soils in different proportions of soft rock and sand. CK: the volume ratio of soft rock to sand is 0:1; C1: the volume ratio of soft rock to sand is 1:5; C2: the volume ratio of soft rock to sand is 1:2; C3: the volume ratio of soft rock to sand is 1:1.

### Cumulative mineralization rate of organic carbon in compound soil

The cumulative mineralization rate of soil organic carbon in different compound ratio of soft rock and sand can reflect the strength of the carbon fixation capacity of the new compound soil. The higher the ratio, the weaker the carbon sequestration capacity of the soil, and vice versa. It can be seen from **Fig 4** that the cumulative mineralization rate of soil organic carbon in the three treatments of CK, C1 and C2 did not reach significant difference after 30 days of culture (P > 0.05), but the soil organic carbon accumulation rate of C1 treated was the lowest in the three treatments. The cumulative mineralization rate of organic carbon treated by C3 was 18.23%, which was significantly increased by 4.62, 6.03 and 5.96 percentage points respectively compared with CK, C1 and C2. Compared with C1 treatment, the cumulative mineralization rate of organic carbon in CK, C2 and C3 treatments increased by 1.41, 0.07 and 6.03 percentage points, respectively.

**Figure 4.**
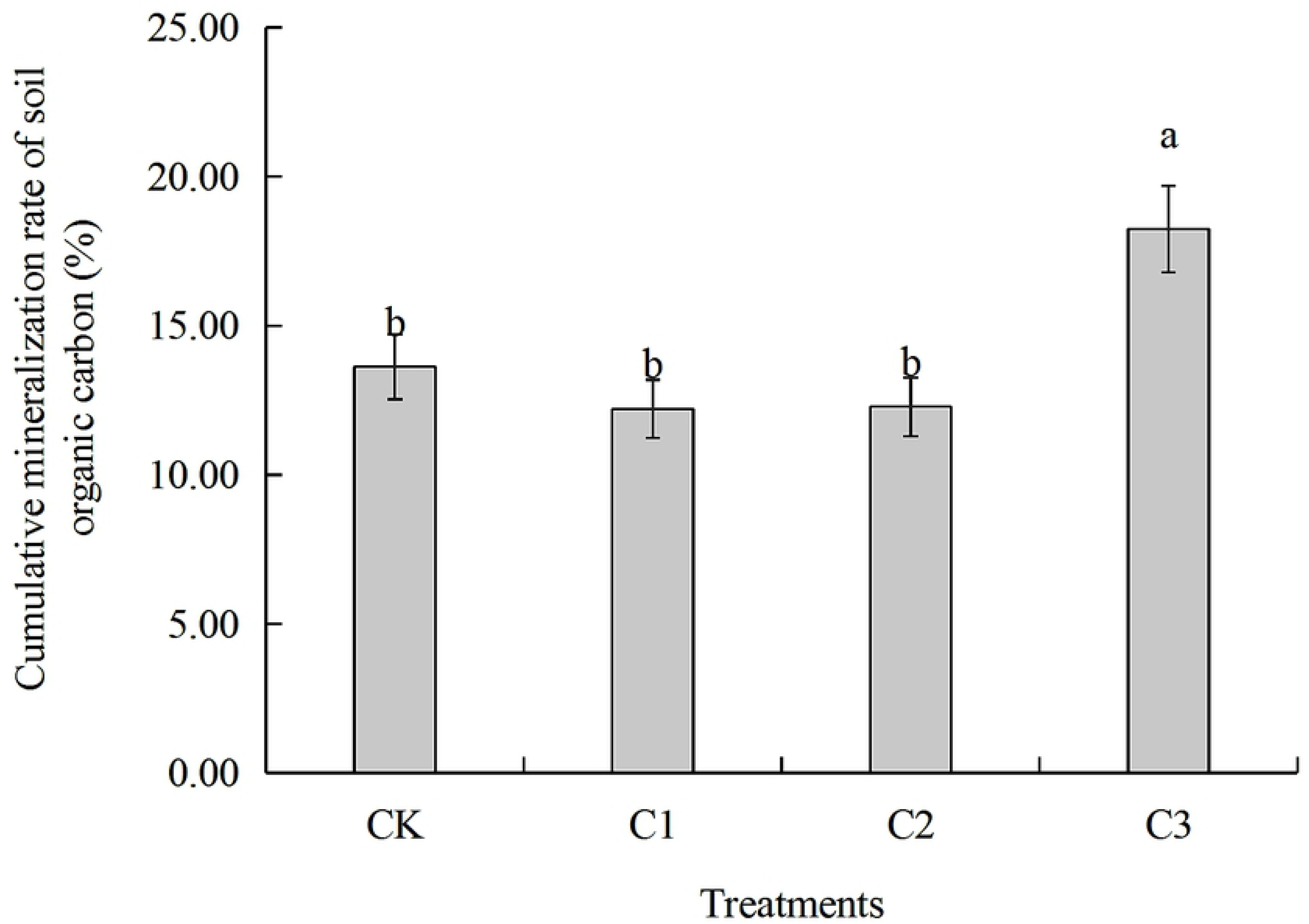
Cumulative mineralization rate of soil organic carbon under different compound ratios during the 30 days of incubation. CK: the volume ratio of soft rock to sand is 0:1; C1: the volume ratio of soft rock to sand is 1:5; C2: the volume ratio of soft rock to sand is 1:2; C3: the volume ratio of soft rock to sand is 1:1. Different letters above the bars mean significant difference (at 0.05 level) between treatments.

### Fitting parameters of organic carbon mineralization in compound soil

There were significant differences between the parameters of the kinetic equations of organic carbon mineralization in the soil of different proportions of soft rock and sand, and the first-order kinetic equation C_t_= C_0_ (1-e^-kt^) was used for parameter fitting (P < 0.01). The potential mineralizable organic carbon (C_0_) content of CK treatment was 257.44 mg kg^-1^, and the C_0_ values in C1, C2 and C3 treatment were significantly increased by 104.34%, 88.24% and 263.95%, respectively (P < 0.05). There was no significant difference between C1 and C2 treatments (**Table 2 and S1 Table**), and C3 treatment significantly increased C_0_ values by 78.11% and 93.34% compared with C1 and C2 treatments. The k values of C1, C2 and C3 treatments were significantly lower than those of CK treatments and there was no significant difference between them. The trend of T_1/2_ was opposite to that of k, indicating that the addition of different proportion of soft rock reduced the mineralization rate constant of organic carbon and increased the retention time of organic carbon in soil. The C_0_/SOC of CK treatment was 12.78%, C1, C2 and CK treatment were not significantly different, C3 treatment was significantly increased by 8.48% compared with CK treatment.

### Compound soil microstructure

Using scanning electron microscope (SEM) to observe 1000 times magnified images of the compound soil, it can be found that from the appearance of individual soil particles, the shape of sand grains was irregular, but the degree of grinding was high, and there are no sharp edges and angles in CK treatment (**Fig 5-a**). With the increase of the volume fraction of the soft rock (C1, C2, C3) (**Fig 5-b, c, d**), the structure of the composite soil has no obvious change, but the filling of fine particles increases gradually. As the volume fraction of small particles increases gradually, the distance between small particles and large particles was larger than that between large particles and large particles. When the content of soft rock reached 50% (C3, **Fig 5-d**), due to the increase of the specific surface area of the small particles, the large particles were not enough to support the soil structure, which shows the collapse of the soil mass.

**Figure 5.**
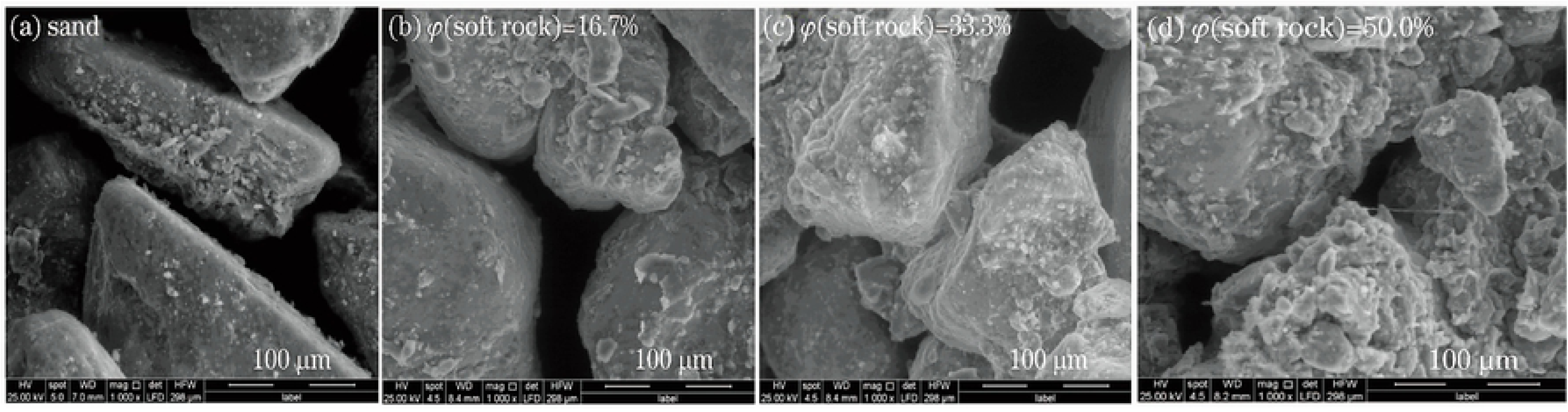
The microstructure of compound soil in different proportions of soft rock and sand. (a): the volume ratio of soft rock to sand is 0:1 (CK); (b): the volume ratio of soft rock to sand is 1:5 (C1); (c): the volume ratio of soft rock to sand is 1:2 (C2); (d): the volume ratio of soft rock to sand is 1:1 (C3).

### Compound soil mechanical composition

In the CK treatment, the content of sand was 86.60%. With the increase of the volume fraction of soft rock, the content of sand gradually decreased, among which C1, C2 and C3 treatment were 33.34, 42.48 and 46.40 percentage points lower than CK treatment respectively (P < 0.05). The silt content increased gradually with the increase of soft rock, and the increase of C1, C2 and C3 was 28.88, 35.48 and 39.16, respectively (P < 0.05). The clay content of CK treatment was 2.28%. With the increase of volume fraction of soft rock, C1, C2 and C3 treatments increased by 4.46, 7.00 and 7.24 percentage points respectively compared with CK treatment (P < 0.05), and the increase of clay component was smaller than that of silt. There were no significant differences between the three treatments of C1, C2 and C3. On the whole, with the increased of the volume fraction of soft rock, the texture of compound soil gradually changed from sandy soil to sandy loam, loam and silty loam.

## Discussion

Soil organic carbon mineralization is an important biochemical process in soil, which is directly related to the release and supply of soil nutrient elements, CO_2_ gas emissions and soil quality maintenance [17]. CK treatment and C3 in the whole culture period with the same mineralization reaction characteristics, reaching a peak on the 3rd day, followed by a rapid and slow decline (**Fig 2**), because the soil microenvironment was still at the beginning of the reaction, it was also possible that the compound soil organic carbon in the initial stage of mineralization was mostly in the form of complex compounds, and there were few small molecular compounds that were easily decomposed. At this time, the microorganism needs to simplify the complex compound before it can be absorbed and utilized, so the respiratory rate showed a rapid rising phase in the initial stage [18]. The reaction characteristics of C1 treatment and C2 treatment were the same, and the trend of decrease was observed throughout the culture period. All the treatments can be divided into two stages: rapid decline on day 1-11 and steady decline on day 11-30. The cumulative mineralization on day 11 reached 54.96%-74.44% of the total mineralization (**Fig 3**), which was consistent with the study of Zhang et al. [19]. The reason was that in the early mineralization stage, the organic matter mainly decomposed by soil microbes is derived from animal and plant residues and their secretions. At this time, there were a large number of active organic substances such as sugars and proteins which were easily decomposed in the soil, which provided abundant carbon sources and nutrients for soil microorganisms and promote microbial activity. With the prolongation of culture time, the active organic components which were easily to decompose in soil were gradually used by the microorganisms, and the remaining components such as lignin and cellulose, which were difficult to decompose and be utilized by microorganisms. Therefore, the mineralization rate showed a trend from fast to slow, and the cumulative mineralization showed a cumulative trend of gradual decrease in release intensity [20]. It can be seen that the amount of soil nutrient plays an important role in the organic carbon mineralization process. Kemmitt et al. [21] studied the mineralization process of a small number of microorganisms and found that after fumigation with chloroform reduced the number of microorganisms by 90%, the mineralization rate of organic carbon among all treatments had no significant difference compared with the control treatment. Kemmitt et al. [21] showed that the mineralization of soil organic matter was limited by a non-biological process that converts the substrate into microbial utilization. At this time, microorganisms plays a small role, and the available organic materials become their limiting factors.

Under the same culture conditions, there were significant differences in soil organic carbon accumulation mineralization (C_t_) with different compounding ratios, which was C3>C1>C2>CK, which showed a consistent trend with the content of organic carbon. (**Fig 1**). The low content of soil organic carbon in CK treatment affects the mineralization rate of soil organic carbon to a certain extent, which was the main reason for the relatively small cumulative release of CO_2_. Aeolian sandy soil has more sand content, larger permeability coefficient and serious water leakage and fertilizer leakage. There are many small particles in soft rock, which are hydrophilic and adsorbent. Mixing soft rock and aeolian sandy soil in a certain proportion can promote the increase of organic carbon content and mineralized amount. The results of this study indicated that the soil clay and silt content increased with the added of the volume fraction of soft rock. When the content of soft rock was 16.7% (C1 treatment), the soil texture was sandy loam (**Fig 6**), and the cumulative mineralization rate of soil organic carbon was also the lowest at this time (**Fig 4**). When the content of soft rock was 33.3% (C2 treatment), the soil texture was loam at this time, and the soil organic carbon accumulation mineralization rate between the C1 treatment and C2 treatment were no significant difference, indicating that the compound ratio of soft rock and sand was 1:5 to 1: 2 can promote the accumulation of soil organic carbon. Because the soft rock content reached 50% (C3 treatment), the large particles was not enough to support the soil structure, showing the collapse of the soil (**Fig 5-d**). At this time, the soil texture was silt loam, and the cumulative mineralization rate was the largest in all treatments. It can be seen that the change of soil structure plays an important role in organic carbon mineralization, and how to change the mechanical properties of soil caused by the change of structure remains to be further studied.

**Figure 6.**
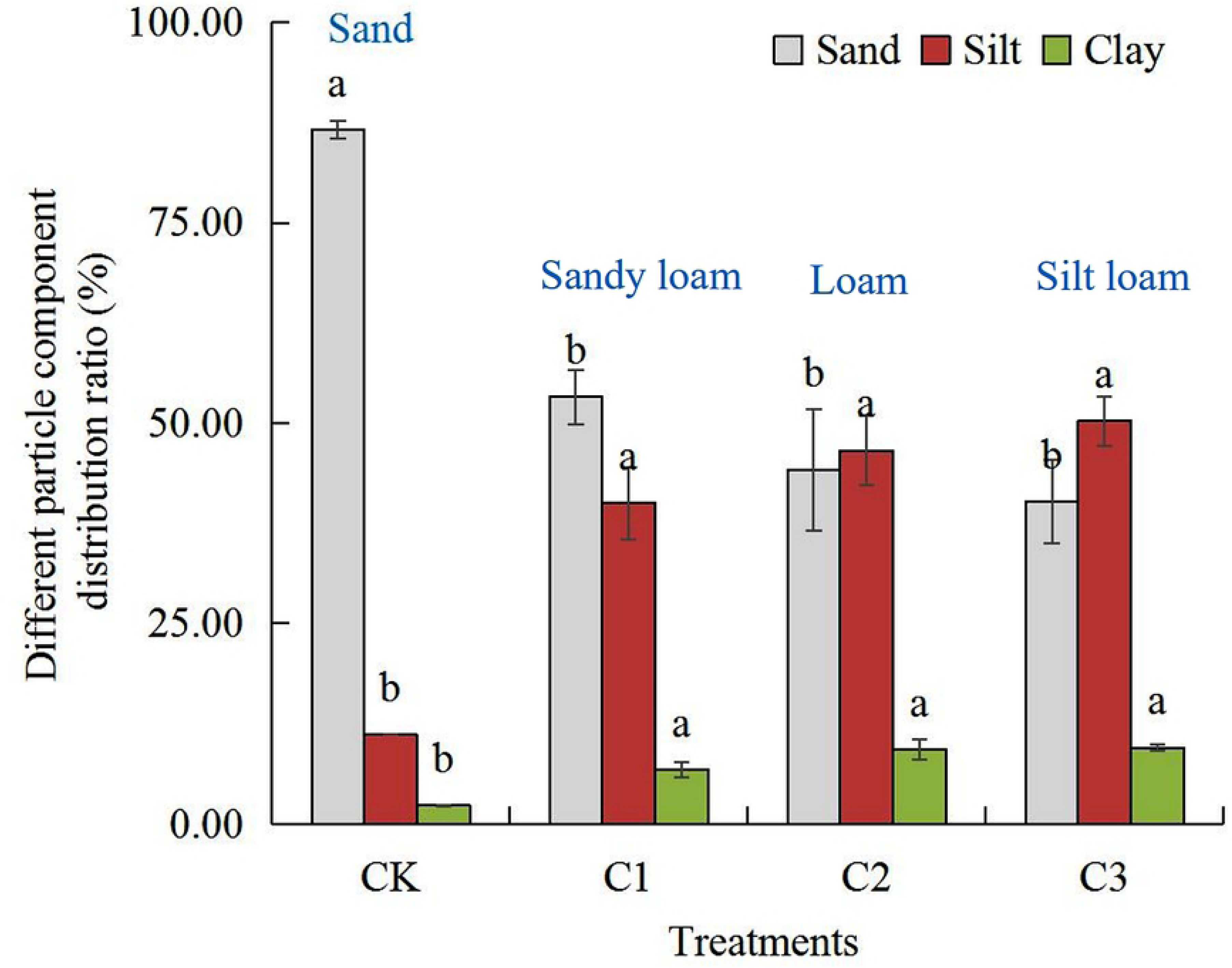
The soil particle composition under different compound ratios of soft rock and sand. CK: the volume ratio of soft rock to sand is 0:1; C1: the volume ratio of soft rock to sand is 1:5; C2: the volume ratio of soft rock to sand is 1:2; C3: the volume ratio of soft rock to sand is 1:1. Lowercase letters indicate significant differences (at 0.05 level) in the same particle composition between treatments.

Soil potentially mineralizable organic carbon (C_0_), also known as biodegradable carbon, is the total amount of organic matter that can be decomposed under the action of microorganisms. The C_0_ value of this study was consistent with the change of C_t_ value, and the specific performance was C3>C1>C2>CK, and there was no significant difference between C1 and C2 treatment. The reason was that with the increased of the content of the soft rock, the non-capillary space between the sand grains was filled by the soft rock, and the capillary pressure increases so as to promote the formation of the soil aggregate structure. The soft rock was also rich in carbonate and other mineral components, as the increased of soft rock volume fraction, the cementation force of the compound soil increased significantly. Because the organic carbon content of the compound soil is significantly higher than that of CK treatment, the activity of plant roots and animals in the compound soil also promotes the fusion of soft rock and sand [22]. Li et al. [23] studied the soil organic carbon mineralization in the Loess Plateau, indicating that the organic carbon mineralization rate constant k was not affected by soil nutrient and pH, but it was related by particle composition. The results of this study showed that there was no significant difference in the value of k between C1, C2 and C3 treatments, and they were significantly lower than CK treatment, and the changes in T_1/2_ and k values were opposited. It may be because the long-term application of chemical fertilizer in this experiment increases the inorganic nitrogen content such as soil nitrate nitrogen and ammonium nitrogen, and reacted with compounds such as lignin or phenol present in the soil, so that organic carbon has lower decomposition property [24, 25]. In addition, it has been reported that the increased of soft rock can promote the formation of aggregates in the compound soil, so that some organic carbon particles can be encapsulated by the aggregates to avoid degradation, thus increasing the retention time of organic carbon in the soil [26,27]. The C_0_/SOC value can reflect the solid storage capacity of the compound soil organic carbon, the larger the ratio, the stronger the soil organic carbon mineralization ability, and vice versa. The results of this study indicated that the C_0_/SOC values in all treatments were C3>CK>C2>C1, and there was no significant difference between C1 and C2, which was consistent with the trend of soil organic carbon accumulation mineralization rate.

## Conclusion

The soil organic carbon content can be significantly increased by the different compound ratios of soft rock and sand. With the increase of the content of soft rock, the content of soil sand was gradually decreased, while the content of clay and silt increased gradually, with the largest increase of silt. The soil texture changed from sand to sandy loam, then to loam and silty loam. The results of scanning electron microscopy showed that the specific surface area between large particles and small particles increased with the increase of volume fraction of soft rock and sand. When the volume fraction of soft rock was 50%, the soil structure collapsed. C1 treatment and C2 treatment had the highest mineralization rate on the first day of culture, while CK treatment and C3 treatment reached the maximum on the third day of culture. The whole culture process can be divided into rapid decline of 1-11 days and slow decline of 11-30 days. With the prolongation of culture time, the accumulation intensity of cumulative mineralization of soil organic carbon was gradually reduced. The cumulative mineralization rate of C1 treatment and C2 treatment was the smallest in all treatments, and C_0_/SOC was consistent with its variation rule. The organic carbon turnover rate was significantly decreased and the retention time in soil was increased with the addition of soft rock, among which C1 treatment and C2 treatment have the best effect. It can be seen that the accumulation of compound soil organic carbon can be significantly increased when the ratio of soft rock to sand was 1:5 to 1:2, which can be used as an important basis for soil remediation measures in farmland.

## Supporting information

**S1 Table. Organic carbon content, organic carbon mineralization rate and mineralization amount, fitting parameters and texture of compound soil.**

(XLS)

## Acknowledgments

We thank research staffs for their contributions to this work.

## Author Contributions

**Conceptualization:** Jichang Han.

**Data curation:** Lei Ge, Chang Tian.

**Formal analysis:** Yan Xu.

**Funding acquisition:** Jichang Han.

**Investigation:** Chendi Shi.

**Methodology:** Zhen Guo.

**Project administration:** Juan Li.

**Resources:** Tingting Cao.

**Supervision:** Yan Xu.

**Writing-original draft:** Zhen Guo.

**Writing-review & editing:** Zhen Guo, Tingting Cao.

